# Tool-sensed object information effectively supports vision for multisensory grasping

**DOI:** 10.1101/2023.01.03.522671

**Authors:** Ivan Camponogara, Alessandro Farnè, Robert Volcic

**Affiliations:** Division of Science, New York University Abu Dhabi, Abu Dhabi, United Arab Emirates; Integrative Multisensory Perception Action and Cognition Team - ImpAct, Lyon Neuroscience Research Center, INSERM U1028, CNRS U5292, Bron, France; UCBL, University of Lyon 1, Villeurbanne, France; Neuro-immersion, Hospices Civils de Lyon, Bron, France; Center for Artificial Intelligence and Robotics, New York University Abu Dhabi, Abu Dhabi, United Arab Emirates

**Keywords:** Multisensory integration, grasping, tool sensing, haptics, vision, tool use

## Abstract

Tools enable humans to extend their sensing abilities beyond the natural limits of their hands, allowing them to sense objects as if they were using their hands directly. The similarities between tool-mediated and hand-based sensing entail the existence of comparable processes for integrating tool- and hand-sensed information with vision, raising the intriguing question of whether tools can support vision in bimanual object manipulations. Here we investigated this question by measuring participants’ performance while reaching for and grasping objects either held with a tool or with their hand. We found that tool-mediated sensing effectively supports vision in multisensory grasping. Even more intriguingly, tool-mediated sensing resembled hand-based sensing. In addition, by manipulating the object features (availability of position and size versus position only), we found that both tool- and hand-mediated action performance was not hindered by the absence of size information. Thus, integrating the tool-sensed position of the object with its vision is sufficient to promote a multisensory advantage in grasping. In sum, our findings indicate that multisensory integration mechanisms significantly improve grasping actions, fine-tuning contralateral hand movements even when object information is only indirectly sensed through the hand operating a tool.

**Significance statement:** Tools allow extending the hands sensing capabilities beyond their anatomical limits. Here we show that object information sensed through a tool can guide bimanual object manipulations as effectively as when directly sensed by the hand. Both tool and hand sensing provide relevant object positional information that are merged with vision to improve action performance. Our findings provide evidence about the interchangeable use of tools and hands for skilled actions and open new perspectives for prosthetic applications and rehabilitative plans.

## Introduction

Evolutionary speaking, tool use is a developmental milestone that allows different species to expand their motor repertoire and facilitate the interactions with objects otherwise unreachable, thus enhancing the chances of survival (Biro, Haslam, & Rutz, 2013). Humans can use tools, such as a grabber, as an extension of the hand to manipulate objects located within and beyond their natural reaching capabilities (Bell & Macuga, 2022; Canzoneri et al., 2013; Cardinali, Brozzoli, Finos, Roy, & Farnè, 2016; Cardinali et al., 2009, 2012; Costantini, Ambrosini, Sinigaglia, & Gallese, 2011; Farnè, Bonifazi, & Làdavas, 2005; Farnè, Iriki, & Làdavas, 2005; Farnè & Làdavas, 2000; Gentilucci, Roy, & Stefanini, 2004; Martel et al., 2019; Martel, Finos, Koun, Farnè, & Roy, 2021; Miller, Cawley-Bennett, Longo, & Saygin, 2017; Sposito, Bolognini, Vallar, & Maravita, 2012) to the same degree as a natural hand grasp at both the kinematic (Gentilucci et al., 2004; Itaguchi & Fukuzawa, 2014) and neural levels (Gallivan, McLean, Valyear, & Culham, 2013; Jacobs, Danielmeier, & Frey, 2010; Johnson-Frey, 2004; Maravita & Iriki, 2004), suggesting common motor and neural control mechanisms for tool and hand mediated object manipulations.

Tools do not only expand our motor capabilities, but they also allow humans to broaden their hand’s inner haptic (proprioceptive and tactile) sensory abilities beyond its anatomical limits (Arbib, Bonaiuto, Jacobs, & Frey, 2009; Burton, 1993; Kilteni & Ehrsson, 2017; Yamamoto & Kitazawa, 2001) to an almost indistinguishable extent from hand-based sensing (Kilteni & Ehrsson, 2017; Miller et al., 2019, 2018; Takahashi, Diedrichsen, & Watt, 2009; Takahashi & Watt, 2014, 2017). For instance, by means of a rod, humans can easily define an object’s position encoding, via haptics, the vibratory patterns elicited by the impact of the rod with the object (Miller et al., 2019, 2018). Concurrently, tools, such as pliers, can be used to detect the size of an object by decoding the distance between the digits holding the pliers (Takahashi et al., 2009; Takahashi & Watt, 2014, 2017). The striking similarity between tool-mediated and hand-based sensing entails a comparable integration process of tool and hand-sensed information with vision. When a tool-held object is also simultaneously seen, the tool-sensed information integrates with vision in a haptic-like manner (Holmes, Sanabria, Calvert, & Spence, 2007; Takahashi et al., 2009; Takahashi & Watt, 2014, 2017), mimicking the same multisensory integration process occurring when the object is sensed by the hand (Ernst & Banks, 2002; Ernst & Bülthoff, 2004; Soto-Faraco, Ronald, & Spence, 2004). Additionally, it has been proposed that tool-mediated and hand-based sensing abilities share a common neural mechanism governing both sensing modalities (Miller et al., 2019). Thus, tools are incredible means to expand motor and sensory human capabilities, resembling hand-like interactions with surrounding objects.

The stunning resemblance between the hand and the tool at the kinematic, perceptual, and neural levels raises the intriguing question of whether tools could support also bimanual object manipulations. In everyday life, we often interact with objects we already hold in our hand (e.g., passing our smartphone from one hand to the other). In this case, the haptic information stemming from hand-based sensing is sufficient to define the main features of an object, such as its size and position, and guide the contralateral hand (Camponogara & Volcic, 2019a, 2019b, 2021, 2022) or tool grasping (Martel et al., 2019). However, when the object is concurrently felt and seen, the integration of redundant haptic ad visual sensory information leads to a superior grasping performance compared to when either vision- or haptic-only inputs are available (Camponogara & Volcic, 2019a, 2019b, 2021, 2022; Pettypiece, Goodale, & Culham, 2010). Within this multisensory-motor integration process, haptics plays a major role in providing positional information (Camponogara & Volcic, 2021, 2022). Thus, it is well established that haptic inputs from the hand holding the object actively support vision in planning and executing accurate grasps. Yet, whether tool-mediated sensing also supports vision for multisensory grasping remains unknown.

Here we filled this gap by examining the grasping performance toward seen and tool- or hand-held objects. In experiment 1, we investigated whether tool-mediated sensing (sensing the to-be-grasped object with a grabber) could support vision in guiding contralateral hand grasping by comparing action performance toward objects that could only be seen (visual condition) or toward seen and tool-held objects (visuo-tool condition). The level of multisensory advantage provided by the tool was further investigated by comparing these conditions with a condition in which objects were simultaneously seen and held by the hand (visuo-haptic condition). If tool-sensed information is irrelevant, we should see an unvaried grasping performance either with or without the additional support of the tool (i.e., similar performance in visuo-tool and visual conditions). In contrast, if tool-mediated sensing supports vision, we expect a superior grasping performance when tool-sensed information is available. Concurrently, if tool-mediated sensing and hand-based sensing support vision in equivalent ways (i.e., same multisensory advantage), we expect grasping actions to be similar in visuo-haptic and visuo-tool conditions. In experiment 2, to exclude that the improvements in grasping kinematics were a mere effect of the force exerted by the hand clenching the tool, a phenomenon known as motor overflow (Addamo, Farrow, Hoy, Bradshaw, & Georgiou-Karistianis, 2007), we compared grasping performance with the tool either directly grasping the object (by exerting force to close the gripper) or simply touching it (without the need to exert any clenching force). If motor overflow plays a role, we expect any tool-mediated sensing advantage to disappear when the object is only touched and not held by the tool. Lastly, in experiment 3, we investigated which tool-sensed information (object size or its position) is used in visuo-tool grasping. According to our previous works on visuo-haptic grasping (Camponogara & Volcic, 2021, 2022), we hypothesized that the tool supports vision by providing mainly positional object information.

## Experiment 1

### Methods

#### Participants

Twenty right-handed participants participated in this experiment (6 males, age 20.2 ± 1.6 years). All had normal or corrected-to-normal vision and no known history of neurological disorders. All of the participants were naïve to the purpose of the experiment and were provided with a subsistence allowance. The experiment was undertaken with the understanding and informed written consent of each participant and the experimental procedures were approved by the Institutional Review Board of New York University Abu Dhabi.

#### Apparatus

The set of stimuli consisted of three 150 mm high 3D-printed cylinders with diameters of 30, 40, 50 mm positioned at 350 mm from the participants’ position. The tool consisted of a 55 cm long claw grabber, whose gripper closed by applying a pressure on its handle (Figure 1a). A 5 mm high rubber bump with a diameter of 9 mm was attached just in front of the participants, 300 mm to the right. This bump was marking the start positions for the right hand. A pair of occlusion goggles (Red Scientific, Salt Lake City, UT, USA) was used to prevent vision of the workspace between trials. A pure tone of 1000 Hz, 100 ms duration was used to signal the start of the trial, while a tone of 600 Hz of the same duration was used to signal its end. Index, thumb and wrist movements were acquired on-line at 200 Hz with sub-millimeter resolution by using an Optotrak Certus system (Northern Digital Inc., Waterloo, Ontario, Canada). The position of the tip of each digit was calculated during the system calibration phase with respect to two rigid bodies defined by three infrared-emitting diodes attached on each distal phalanx (Nicolini, Fantoni, Mancuso, Volcic, & Domini, 2014). An additional marker was attached on the styloid process of the radius to monitor the movement of the arm. The Optotrak system was controlled by the MOTOM toolbox (Derzsi & Volcic, 2018).

**Figure 1:**
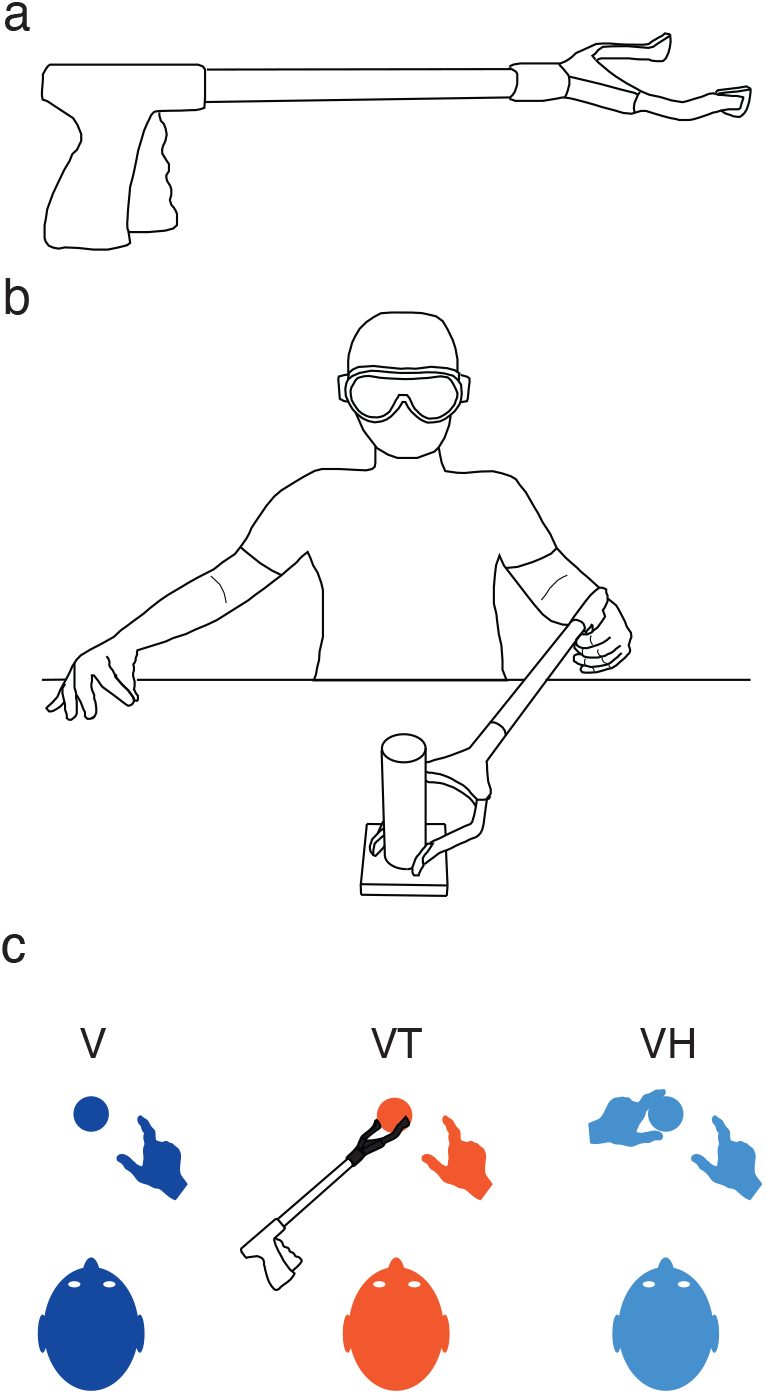
Experiment 1 setup and procedure. a) The tool used in the experiment was a grabber of 55 cm length. The gripper was closed by pressing the handle. b) Experimental setup. Participant’s sat at the edge of the table with the occlusion goggles on. Grasping actions were always performed with the right hand, and the tool was operated with the left hand. The picture represents the starting position in the visuo-tool condition. c) Representation of the task in each condition (top view). In the V condition, participants had to perform a visually-guided reach-to-grasp movement. In the VT condition, objects were concurrently held with the tool, whereas in the VH condition, objects were held with the left hand.

#### Procedure

Participants sat comfortably at the table with their torsos touching its edge. All the trials started with the participants’ thumb and index digit of the right hand positioned on the start positions, the left hand on the side, either free or with the tool, and the shutter goggles closed. Before each trial, one of the objects was positioned in front of the participant. In the Visual condition (V) the goggles turned transparent, and participants had only visual information about the object. In the Visuo-Tool condition (VT), the experimenter signaled to the participants to move the tool’s gripper to the object and close it on the object’s base. Then, the experimenter triggered the goggles, which turned transparent, and participants were thus able to see and feel the object through the tool. In the Visuo-Haptic condition (VH), the experimenter signaled to the participants to hold the object with their left hand index and thumb at its base (i.e., sense its size and position by means of tactile and proprioceptive inputs). Once both fingers contacted the object, the experimenter triggered the goggles, which turned transparent, allowing participants to have visual and haptic information about the object (Figure 1b). Thus, while in VH, the object size and position were sensed through haptics, in VT these properties were sensed through the tool (Figure 1c). After a variable period (1–1.5 s), the start tone was delivered and participants had to reach for and grasp the object with their right hand. Movements were performed at a natural speed, and no speed constraints were imposed. After 3 s, the end sound was delivered, and participants had to move their right and left hands/tool back to the start positions, and then the goggles turned opaque. Another object was then selected, and the next trial was ready to start. The order of conditions was randomized across participants, while object sizes were randomized within each condition. We ran 15 repetitions for each object size, which led to a total of 135 trials per participant (45 for each condition). Before the experiment, a training session was performed in which ten trials were run in each condition to accustom the participants to the task.

#### Data analysis

Kinematic data were analyzed in R (R Core Team, 2020). The raw data were smoothed and differentiated with a third-order Savitzky-Golay filter with a window size of 21 points. These filtered data were then used to compute velocities and accelerations in three-dimensional space for each digit and wrist. Movement onset was defined as the moment of the lowest, non-repeating wrist acceleration value prior to the continuously increasing wrist acceleration values (Volcic & Domini, 2016), while the end of the grasping movement was defined on the basis of the Multiple Sources of Information method (Schot, Brenner, & Smeets, 2010). We used the criteria that the grip aperture is close to the size of the object, that the grip aperture is decreasing, that the second derivative of the grip aperture is positive, and that the velocities of the wrist, thumb, and index finger are low. Moreover, the probability of a moment being the end of the movement decreased over time to capture the first instance in which the above criteria were met. Trials in which the reaction time was lower than 50 ms or exceeded 900 ms, the end of the movement was not captured correctly or in which the missing marker samples could not be reconstructed using interpolation were discarded from further analysis, the exclusion of these trials (285 trials, 10.5% in total) left us with 2415 trials.

We focused our analyses on three dependent variables: the response time, defined as the time from the start tone to the movement onset, the peak velocity of the hand movement, defined as the highest wrist velocity along the movement, and the peak grip aperture, defined as the maximum Euclidean distance between the thumb and the index finger. We analyzed the data using Bayesian linear mixed-effects models, estimated using the brms package (Bürkner, 2017), which implements Bayesian multilevel models in R using the probabilistic programming language Stan (Carpenter et al., 2017). The models included as fixed-effects (predictors) the categorical variable Condition (V, VH, and VT) in combination with the continuous variable Size. This latter was centered before being entered into the models. Thus, the estimates of the Condition parameters (*β_Condition_*) correspond to the average performance of each Condition. The estimates of the parameter Size (*β_Size_*) correspond instead to the change in the dependent variables as a function of the object size. All models included independent random (group-level) effects for subjects. Models were fitted considering weakly informative prior distributions for each parameter to provide information about their plausible scale. For the response time, we used log-normal priors for the Condition, whereas Gaussian priors for the Size, whereas for the peak velocity and peak grip aperture, we used Gaussian priors for the Condition and Size fixed-effect predictors based on our previous studies (Camponogara & Volcic, 2019a, 2019b, 2021, 2022) (response time *β_Condition_*: median = 5.4 and sd = 0.7, *β_Size_*: mean = 0 and sd = 0.9; peak velocity *β_Condition_*: mean = 1000 and sd = 100, *β_Size_*: mean = 0 and sd = 2; peak grip aperture *β_Condition_*: mean = 85 and sd = 20, *β_Size_*: mean = 0 and sd = 2). For the group-level standard deviation parameters and sigmas we used Student *t*-distribution priors (response time sd parameters and sigma: *df* = 3, scale = 2.5; peak velocity sd parameters and sigma: *df* = 3, scale = 121.5; peak grip aperture sd parameters and sigma: *df* = 3, scale = 12). Finally, we set a prior over the correlation matrix that assumes that smaller correlations are slightly more likely than larger ones (LKJ prior set to 2).

For each model, we ran four Markov chains simultaneously, each for 4,000 iterations (1,000 warm-up samples to tune the MCMC sampler) with the delta parameter set to 0.9 for a total of 12,000 postwarm-up samples. Chain convergence was assessed using the 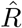 statistic (all values equal to 1) and visual inspection of the chain traces. Additionally, the predictive accuracy of the fitted models was estimated with leave-one-out cross-validation by using the Pareto Smoothed Importance Sampling. All Pareto k values were below 0.5.

The posterior distributions we have obtained represent the probabilities of the parameters conditional on the priors, model and data, and they represent our belief that the “true” parameter lies within some interval with a given probability. We summarize these posterior distributions by computing the medians and the 95% Highest Density Intervals (HDI). The 95% HDI specifies the interval that includes, with a 95% probability, the true value of a specific parameter. To evaluate the differences between parameters of two conditions, we have simply subtracted the posterior distributions of *β_Condition_* and *β_Size_* weights between specific conditions. The resulting distributions are denoted as the credible difference distributions and are again summarized by computing the medians and the 95% HDIs.

For statistical inferences about the *β_Size_* we assessed the overlap of the 95% HDI with zero. A 95% HDI that does not span zero indicates that the predictor has an effect on the dependent variable. For statistical inferences about the differences of the model parameters, *β_Condition_* and *β_Size_*, between conditions, we applied an analogous approach. A 95% HDI of the credible difference distribution that does not span zero is taken as evidence that the model parameters in the two conditions differ from each other.

### Results and Discussion

We found that movements were released earlier (response time) and performed with a narrower peak grip aperture in the visuo-tool compared to the visual condition (Figure 2, panels a, and g), suggesting a successful integration of tool-mediated and visual sensory information. Interestingly, the response time and peak of grip aperture in the visual-tool condition were similar to the visuo-haptic condition. Corroborating our previous results (Camponogara & Volcic, 2019a, 2019b, 2021, 2022), we found a faster action with a narrower peak grip aperture in the visuo-haptic compared to the visual condition.

**Figure 2:**
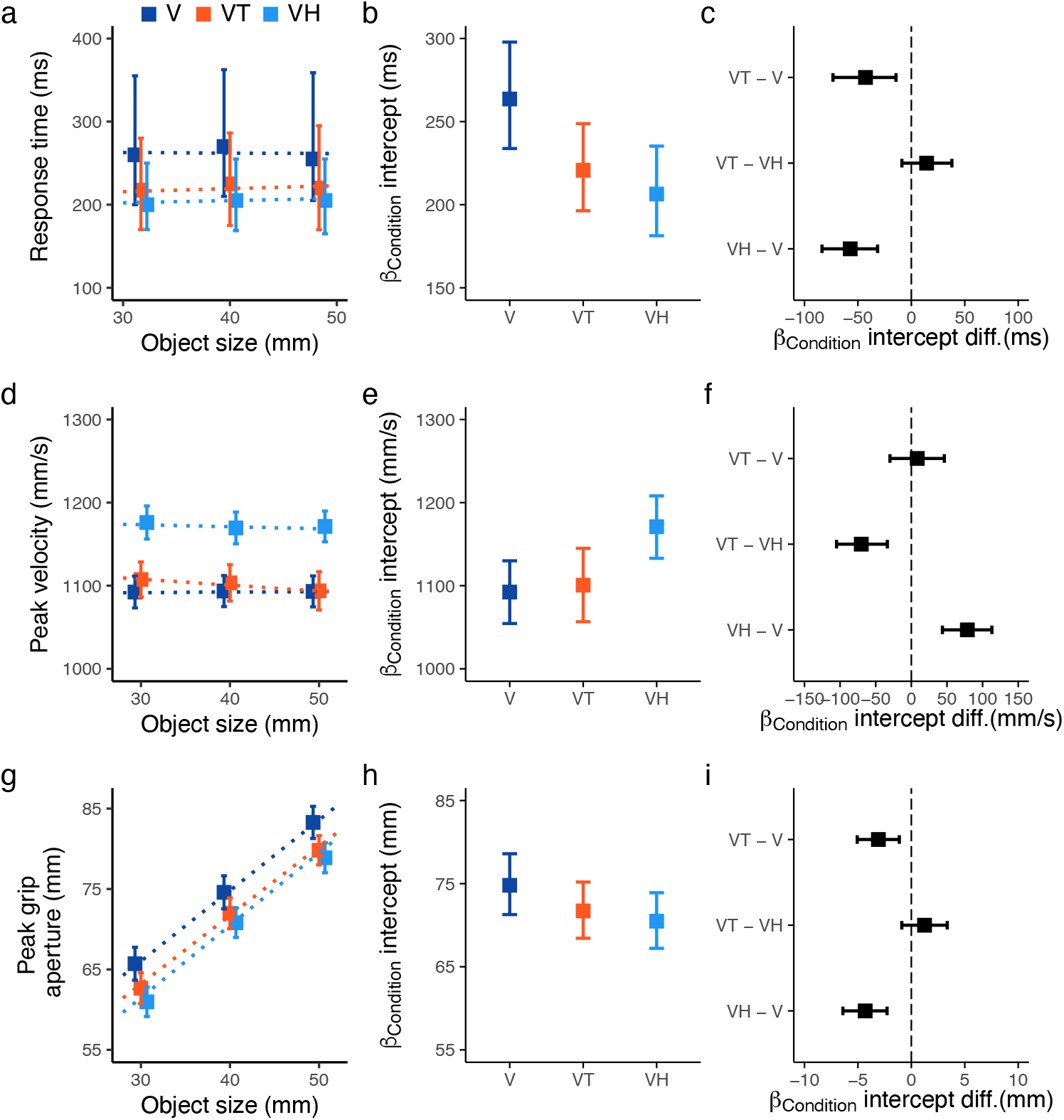
Summary of Experiment 1 results. Top row: Response time; Middle row: Peak velocity; Bottom row: Peak grip aperture results. a) Median, d, g) Data averaged as a function of the object size, b, e, h) Posterior beta weights of the Bayesian linear mixed-effects regression model for the predictor Condition, c, f, i) Credible difference distributions between conditions for the predictor Condition. In panel a the error bars represent the interquartile range, in panels d, and g the error bars represent the standard error of the mean. Dotted lines show the Bayesian mixed-effects regression model fits. In panels b, c, e, f, h, i the error bars represent the 95% HDIs of the distributions.

The response time was modulated according to the available sensory information, with an advantage when the object was concurrently seen and held by the tool (VT = 220 ms, 95% HDI = 196 ms, 248 ms) or the hand (VH = 206 ms, 95% HDI = 181 ms, 235 ms) compared to when it was only seen (V = 263 ms, 95% HDI = 233 ms, 297 ms). The response time was credibly lower in VT compared to V and in VH compared to V, with no differences between VT and VH (Figure 2, panels b and c). The response time was not affected by changes in object size in any of the conditions, with slope values ranging between −0.25 and 0.15 corresponding to minimal variations in response time between the smallest and the largest object (~5 ms difference equivalent to ~2% of the average response time).

The peak velocity was modulated according to the available sensory information as well, but actions were equally fast when the object was only seen, or both seen and held with the tool (Figure 2 panels e and f). We replicated our previous finding by showing a credibly higher peak velocity in VH compared to V (Camponogara & Volcic, 2019b, 2021, 2022). The peak velocity was similar between V and VT, whereas it was also credibly higher in VH compared to VT. Additionally, it was not affected by a change in the object size in V and VH, and showed only a minimal variation in VT. Results showed a slope value in VT of −0.7 (HDI: −1.23, −0.20), which corresponds to a change of ~14 mm/s from the largest to the smallest objects (equivalent to ~1% of the average peak velocity)

The peak grip aperture was also clearly affected by the available sensory inputs (Figure 2 panels h and i). The peak grip aperture was smaller in VT compared to V (VT = 72 mm, 95% HDI = 68 mm, 75 mm), and similar between VT and VH conditions. These patterns of results were remarkably consistent across participants (Figure 3), suggesting that position and size information sensed from the tool can be effectively used to aid vision and improve grasping performance. Peak grip aperture was credibly smaller in the VH condition compared to the V condition (VH = 70 mm, 95% HDI = 67 mm, 73 mm;V = 75 mm, 95% HDI = 71 mm, 78 mm), confirming that the simultaneous availability of visual and haptic inputs leads to a multisensory advantage (Camponogara & Volcic, 2019b, 2021, 2022).

**Figure 3:**
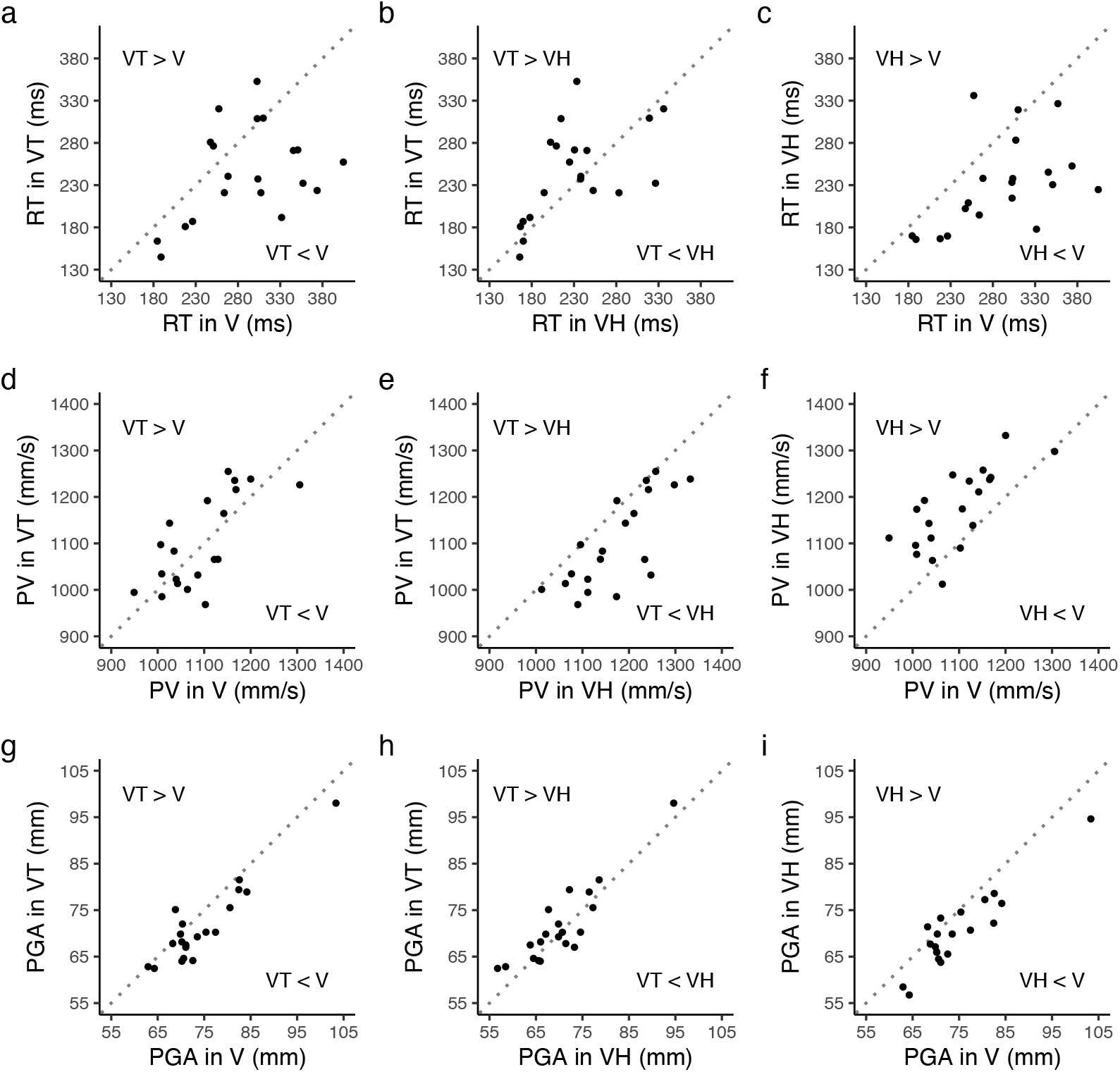
Scatterplots of paired observations in Experiment 1. Each point represents the median response time (top row, RT), average peak velocity (middle row, PV), and average peak of grip aperture (bottom row, PGA) of a single participant for a pair of conditions: first column (a, d, g) VT and V, second column (b, e, h) VT and VH, third column (c, f, i) VH and V. The diagonal reference line of no effect has slope 1 and intercept 0. Points above the diagonal line indicate that the variable of the condition represented on the ordinate axis is larger than the variable represented on the abscissa.

Thus, tool-sensed object information was effectively integrated with vision to support grasping performance. Tool-mediated sensing may have occurred by translating the haptic information from the hand holding the tool in positional and size information. Specifically, a change in the object size required modulation of the clenching force exerted on the handle to close the tool’s gripper on the object. This force modulation was also associated with a concurrent change of the distance between the thumb and the other four fingers, a cue that may have been used to infer the object size (Berryman, Yau, & Hsiao, 2006; Takahashi et al., 2009; Takahashi & Watt, 2014). Concurrently, the pattern of somatosensory inputs generated by the impact of the tool’s gripper with the object (Miller et al., 2019, 2018) and the haptic inputs stemming from the inertia generated by the active placement of the tool’s gripper on the object (Chan, 1994; Solomon & Turvey, 1988; Solomon, Turvey, & Burton, 1989) may have been used to infer the tool’s length and, consequently, the object’s position. However, the faster action initiation and the smaller peak grip aperture observed when vision was complemented with tool-sensed information may have been alternatively aroused from the influence of the clenching force exerted on the tool’s handle. Studies on bimanual interactions showed that exerting force with one limb generates a mirrored involuntary movement of the homologous muscles in the contralateral limb, a phenomenon known as “motor overflow” (see Addamo et al. (2007) for a review). Even though its origin is still under debate, several studies claim that the motor overflow stems from a facilitatory effect of the movement-related cortical regions onto the homologous contralateral areas in the opposite hemisphere (Hoy, Fitzgerald, Bradshaw, Armatas, & Georgiou-Karistianis, 2004). Thus, exerting a clenching force on the tool’s handle might have promoted a pre-activation of the cortical areas involved in the contralateral limb control, which enabled a faster release of the motor plan with a concurrent reduction of the overall grip aperture. To test whether the results in the VT condition were due to the motor overflow, a second experiment was performed where we manipulated the clenching force exerted on the handle by asking participants to either close or not the tool’s gripper on the to-be-grasped object. If actions are affected by the motor overflow, we expected the right-hand reach-to-grasp actions to be released earlier (i.e., shorter response time), and with a smaller grip aperture when the gripper is closed than when it is not.

## Experiment 2

### Methods

#### Participants

Nineteen right-handed new participants took part in Experiment 2 (5 males, age 20.9 ± 2.9 years). All had normal or corrected-to-normal vision and no known history of neurological disorders. All of the participants were naïve to the purpose of the experiment and were provided with a subsistence allowance. The experiment was undertaken with the understanding and informed written consent of each participant, and the experimental procedures were approved by the Institutional Review Board of New York University Abu Dhabi.

#### Apparatus

The experimental setup was the same as in Experiment 1, except that a new set of stimuli was used, which consisted of three cylinders of 60 mm height supported by a 60 mm high cylindrical post of 10 mm diameter (Figure 4a). The upper part of these stimuli was identical to the first set of stimuli and thus varied in diameter across trials. The post supporting the upper part had instead a fixed diameter. Thus, enclosing the gripper on the post led to a constant clenching force level on the handle across the different object sizes.

**Figure 4:**
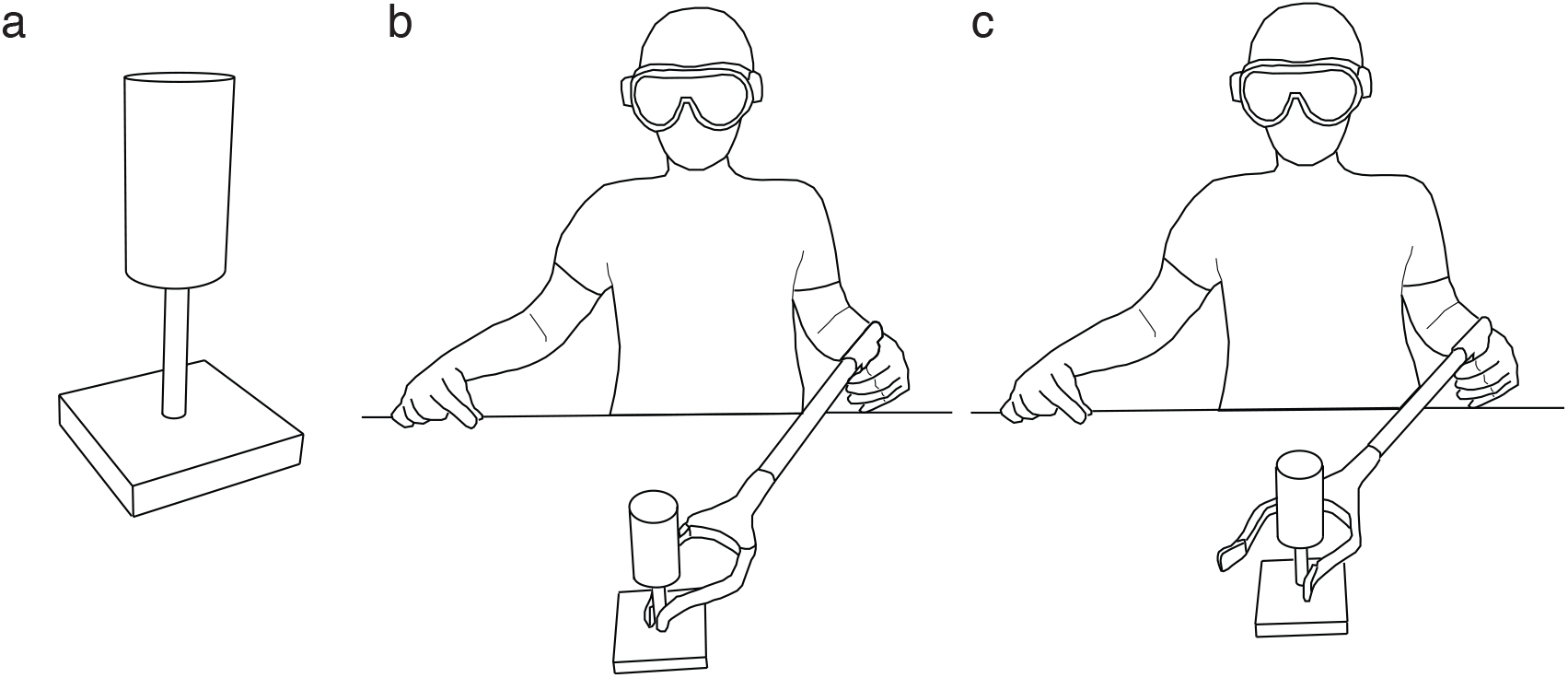
Experiment 1 setup and procedure. a) Example of a stimulus used in Experiment 2. b) Representation of the task in the Visuo-Tool-Closed condition. Participants closed the tool’s gripper on the post supporting the to-be grasped object c) Representation of the task in the Visuo-Tool-Open condition. Participants kept the tool’s gripper open, with the left tip of the gripper touching the post supporting the object.

#### Procedure

The procedure was the same as for the Visuo-Tool condition of Experiment 1. In a Visuo-Tool-Closed condition (VTC) participants closed the tool’s gripper around the post supporting the object (Figure 4b), whereas in the Visuo-Tool-Open condition (VTO) the gripper was kept open, with the left tip in contact with the object’s post (Figure 4c). The order of the conditions was randomized across participants. Object sizes were randomized within each condition and 15 trials were performed for each object size and condition, which led to a total of 90 trials per participant. Before each condition, participants underwent a training session of ten trials to get accustomed with the task.

#### Data analysis

The raw data processing and the statistical analyses were identical to those of Experiment 1. Based on the same exclusion criteria, a total of 319 trials (~17% in total) were excluded which left us with 1481 trials for the final analysis. As in Experiment 1, we focused our analyses on the reaction time, peak velocity and peak grip aperture. The 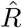 statistic and visual inspection of the chain traces confirmed successful chains convergence. All Pareto k values were below 0.5. As in Experiment 1, we report the posterior distribution of the *β_Condition_* and *β_Size_* for each condition, and contrast the different conditions by computing the differences between the posterior distributions for each predictor.

### Results and Discussion

Movements were almost indistinguishable between the two conditions, with a slight higher velocity when the tool’s gripper was closed compared to when it was open (Figure 5).

**Figure 5:**
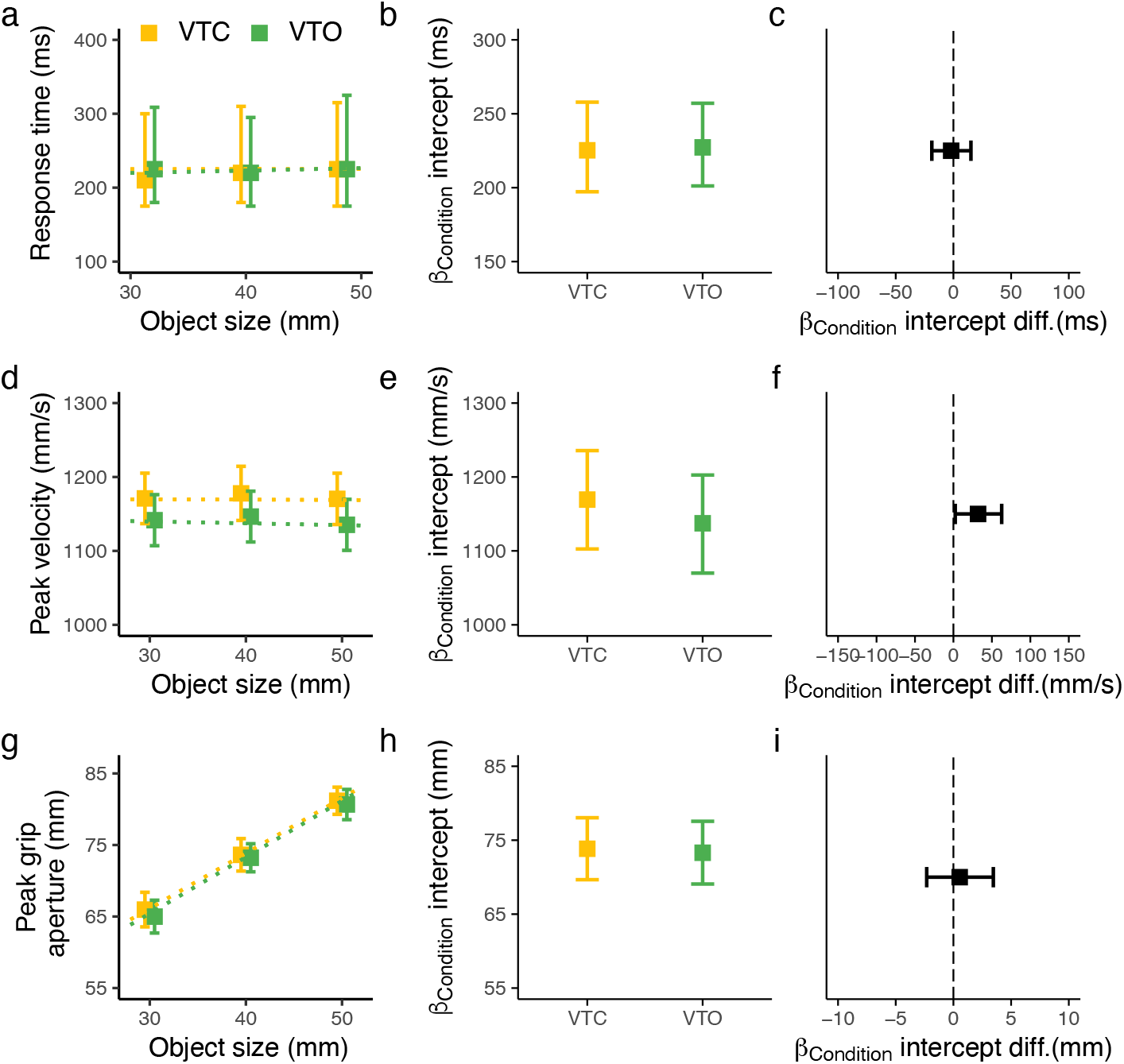
Summary of Experiment 2 results. Top row: Reaction time; Middle row: Peak velocity; Bottom row: Peak grip aperture results. a) Median, d, g) Data averaged as a function of the object size, b, e, h) Posterior beta weights of the Bayesian linear mixed-effects regression model for the predictor Condition, c, f, i) Credible difference distributions between conditions for the predictor Condition. In panel a the error bars represent the interquartile range, in panels d, and g the error bars represent the standard error of the mean. The dotted lines show the Bayesian mixed-effects regression model fits. In panels b, c, e, f, h, i the error bars represent the 95% HDIs of the distributions.

Actions initiated approximately ~220 ms following the start tone for both conditions (VTC = 225 ms, 95% HDI = 197 ms, 257 ms, VTO = 227 ms, 95% HDI = 201 ms, 257 ms), with no modulation according to the object size (slope values of 0.10 and 0.12 for the VTC and VTO conditions respectively). The peak velocity was slightly higher in VTC compared to the VTO condition (mean difference = 32 mm/s, 95% HDI = 1 mm/s, 63 mm/s; VTC = 1169 mm/s, 95% HDI = 1104 mm/s, 1233 mm/s; VTO = 1137 mm/s, 95% HDI = 1074 mm/s, 1201 mm/s), suggesting a slight effect of the motor overflow on the contralateral hand’s reaching, whereas the peak grip aperture and their scaling was identical between the two conditions (VTC = 73 mm, 95% HDI = 69 mm, 78 mm; VTO = 73 mm, 95% HDI = 69 mm, 77 mm). As seen in the Experiment 1, these patterns of results were consistent across participants (Figure 6).

**Figure 6:**
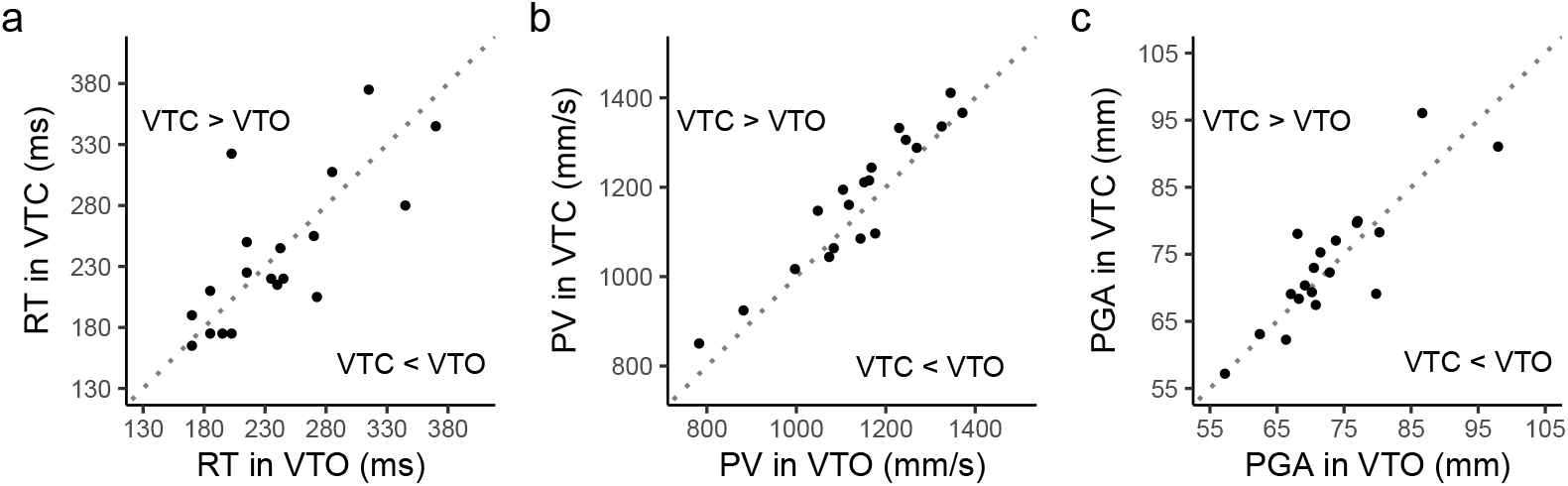
Scatterplots of paired observations in Experiment 2. Each point represents the median response time (a), average peak velocity (b), and average peak of grip aperture (c) of a single participant for a pair of conditions. The diagonal reference line of no effect has slope 1 and intercept 0. Points above the diagonal line indicate that the variable of the condition represented on the ordinate axis is larger than the variable represented on the abscissa.

Taken together our results support that action performance toward a seen and a tool-held object in Experiment 1 resulted from the concurrent use of tool and vision. It’s also interesting to notice that by grabbing the post supporting the object, only positional and not size information could be sensed through the tool. Even though the tool-sensed size was prevented, results were comparable to those found in the VT condition of Experiment 1, where both position and size were available. This suggests that in both experiments, the tool may have supported vision by mainly providing relevant positional information, which was integrated with visual position and size to enable a superior grasping performance. If this is the case, the tool may have played an equivalent role of haptics in visuo-haptic grasping. Previous studies showed that haptic object position is indeed sufficient to produce the typical multisensory advantage characterizing actions toward seen and held objects (Camponogara & Volcic, 2021, 2022). To further investigate whether the tool-sensed positional information is sufficient to promote a grasping advantage, we ran a third experiment where we manipulated the availability of tool-sensed object size information. If the tool-sensed size is crucial for the grasping performance, preventing access to haptic object size will increase the peak of the grip aperture and reduce the grip aperture scaling compared to when the object size is available. In contrast, if the position is sufficient to provide an advantage, we expect a comparable performance either with or without the concurrent presence of size information. According to our previous results with hand-held and seen objects, in either case, we expect no change in the response time and peak velocity since in both conditions, positional information is constantly provided (Camponogara & Volcic, 2021).

## Experiment 3

### Methods

#### Participants

Twenty right-handed new participants took part in Experiment 3 (6 males, age 21.3 ± 2.77). All had normal or corrected-to-normal vision and no known history of neurological disorders. All of the participants were naïve to the purpose of the experiment and were provided with a subsistence allowance. The experiment was undertaken with the understanding and informed written consent of each participant, and the experimental procedures were approved by the Institutional Review Board of New York University Abu Dhabi.

#### Apparatus

We made use of the same experimental setups and sets of objects already used in Experiment 1 and Experiment 2. To recap, the first set of objects consisted in three cylinders whose diameter of 30, 40, 50 mm was constant along their whole height (150 mm). The second sets of objects consisted of three cylinders of 75 mm height supported by a 75 mm high cylindrical post of 10 mm diameter (Figure 7a). The upper part of these stimuli was identical to the first set of stimuli and thus varied in diameter across trials. The post supporting the upper part had a fixed diameter instead.

**Figure 7:**
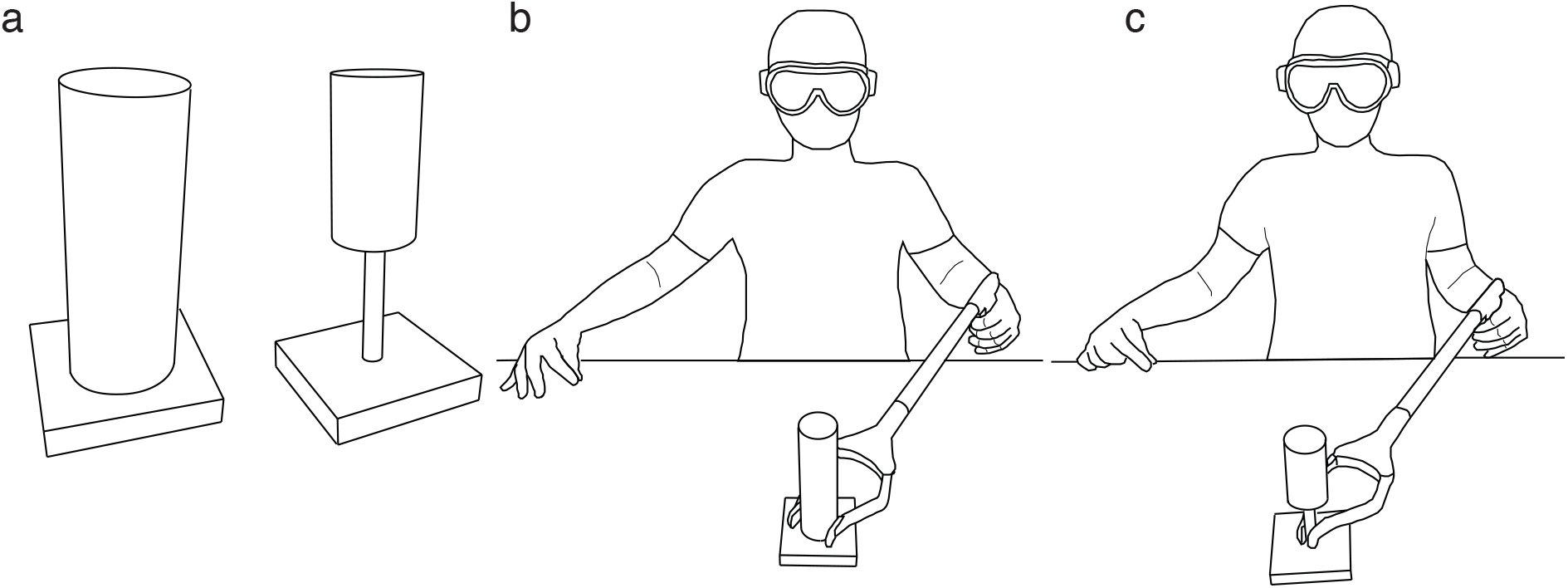
Experiment 1 setup and procedure. a) Example of the sets of objects used in Experiment 3. b) Representation of the task in the Visuo-Tool condition. Participants closed the tool’s gripper on the base of the object, thus sensing both position and size information through the tool c) Representation of the task in the Visuo-Tool-Closed condition. Participants closed the tool’s gripper on the post supporting the to-be grasped object, thus sensing only position information through the tool.

#### Procedure

We chose the VT and the VTC conditions of Experiment 1 and Experiment 2, respectively. In the VT condition participants were presented with the first sets of objects, thus having access to both tool-sensed position and size information (Figure 7b). In the VTC condition the second set of objects was presented; thus, the tool could be used only to sense object positional information (Figure 7c). The order of the conditions was randomized across participants, whereas the object sizes were randomized within each condition. Fifteen trials were performed for each object size and condition, which led to a total of 45 trials per condition (90 per participants). Before each condition, participants underwent a training session of ten trials to get accustomed with the task.

#### Data analysis

The raw data processing and the statistical analyses were identical to those of Experiment 1 and Experiment 2. Based on the same exclusion criteria, a total of 159 trials (~8% in total) were excluded which left us with 1641 trials for the final analysis. As in Experiment 1 and Experiment 2, we focused our analyses on the response time, peak velocity, and peak grip aperture. The 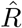 statistic and visual inspection of the chain traces confirmed successful chains convergence. All Pareto k values were below 0.5. We reported the posterior distribution of the *β_Condition_* and *β_Size_* for each condition, and contrast the different conditions by computing the differences between the posterior distributions for each predictor.

### Results and Discussion

Results showed identical movements either with or without the concurrent availability of object size information (Figure 8), and resembled those obtained in the VT and VTC conditions in Experiment 1 and Experiment 2, respectively.

**Figure 8:**
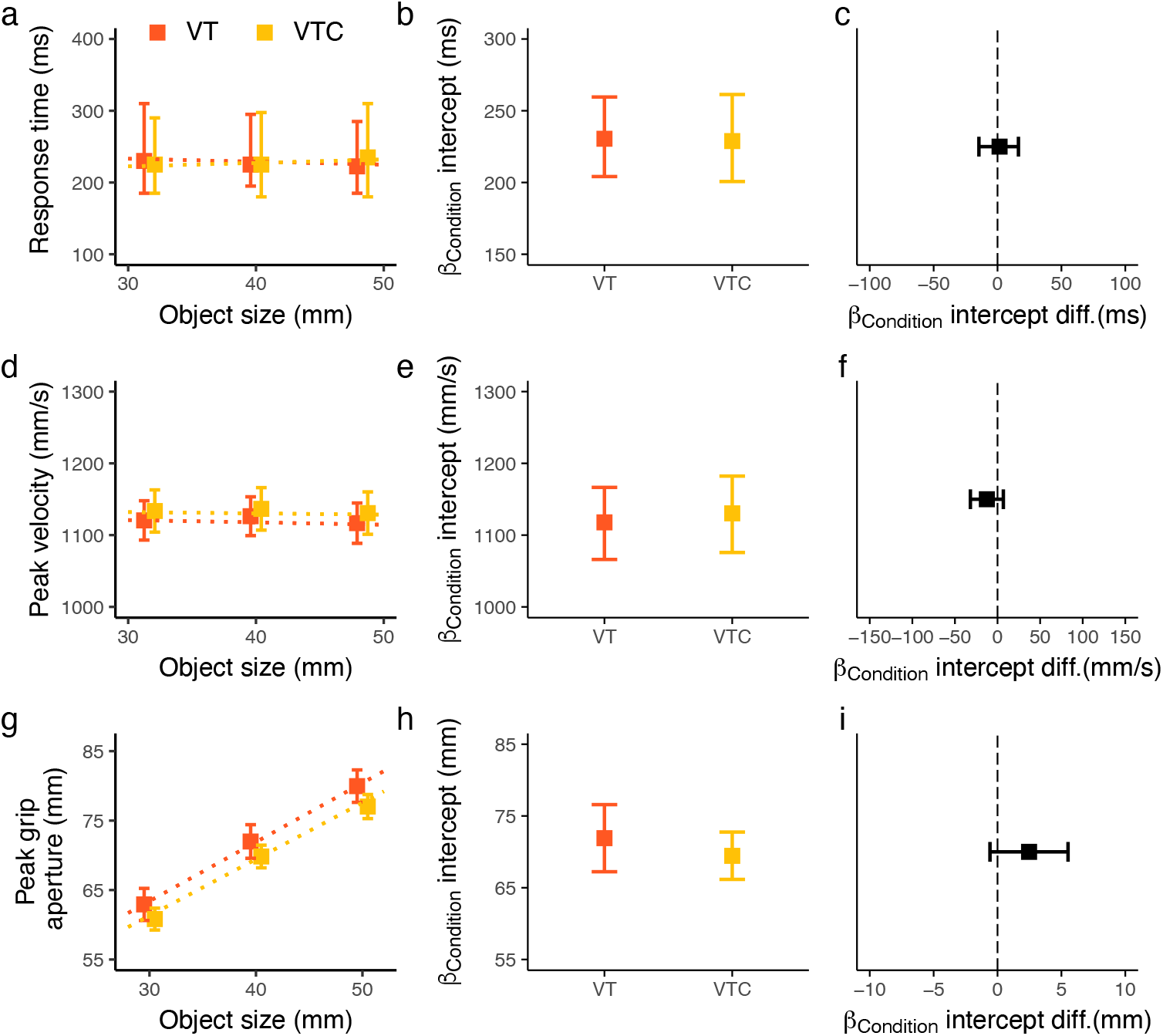
Summary of Experiment 3 results. Top row: Response time; Middle row: Peak velocity; Bottom row: Peak grip aperture results. a) Median, d, g) Data averaged as a function of the object size, b, e, h) Posterior beta weights of the Bayesian linear mixed-effects regression model for the predictor Condition, c, f, i) Credible difference distributions between conditions for the predictor Condition. In panel a the error bars represent the interquartile range, in panels d, and g the error bars represent the standard error of the mean. The dotted lines show the Bayesian mixed-effects regression model fits. In panels b, c, e, f, h, i the error bars represent the 95% HDIs of the distributions.

The action plan was released ~230 ms following the start tone (VT = 230 ms, 95% HDI = 204 ms, 259 ms, VTC = 228 ms, 95% HDI = 200 ms, 261 ms), with no modulation according to the object size (slope values of −0.42 and 0.77 for the VT and VTC conditions respectively). The peak velocity was identical in both conditions (VT = 1118 mm/s, 95% HDI = 1067 mm/s, 1171 mm/s; VTC = 1130 mm/s, 95% HDI = 1073 mm/s, 1187 mm/s), again with no modulation according to the object size (slope values of −0.27 and −0.16 for the VT and VTC conditions respectively). Interestingly, the peak grip aperture and its scaling were indistinguishable between conditions as well (peak grip aperture: VT = 72 mm, 95% HDI = 67 mm, 76 mm; VTC = 69 mm, 95% HDI = 66 mm, 73 mm; scaling peak grip aperture: VT = 0.85 mm, VTC = 0.81 mm), confirming the main role of the tool-sensed position within the sensory integration process. These patterns of results were very consistent across participants (Figure 9).

**Figure 9:**
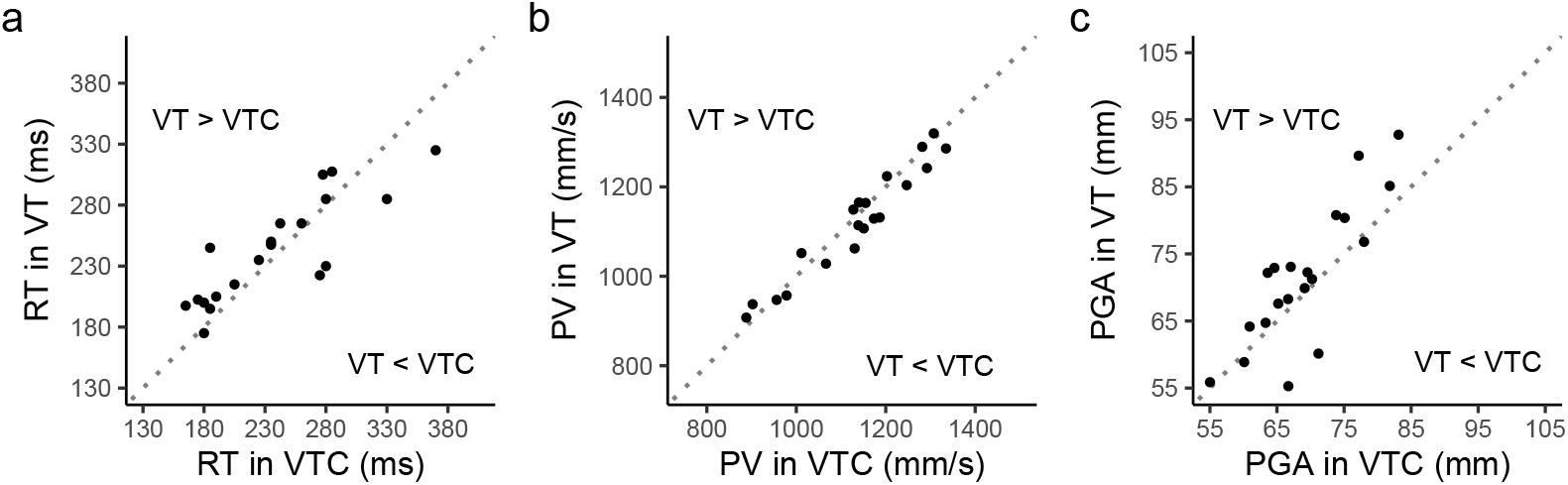
Scatterplots of paired observations in Experiment 3. Each point represents the median reaction time (a), average peak velocity (b) and average peak of grip aperture (c) of a single participant for a pair of conditions. The diagonal reference line of no effect has slope 1 and intercept 0. Points above the diagonal line indicate that the variable of the condition represented on the ordinate axis is larger than the variable represented on the abscissa.

Thus, as seen for visuo-haptic grasping (Camponogara & Volcic, 2021, 2022), results showed that also the tool supports vision by providing mainly positional information. This further corroborates the hypothesis that tools can extend the sensory capacity beyond the body and sensory inputs from the tool can be used as those coming from our own limb (Miller et al., 2019; Miller, Jarto, & Medendorp, 2022; Miller et al., 2018). Here we extended such findings by showing that these tool-sensed information can be actively used to guide a contralateral hand’s grasping.

## General Discussion

In this study, we demonstrated that tool-mediated sensing can guide skilled bimanual object manipulations: humans can successfully integrate tool-mediated object information with vision to guide contralateral hand grasping. Grasping movements were not affected by the clenching force exerted on the handle of the tool, suggesting a genuine combination of tool-sensed information with vision. Even more intriguingly, we found that tool-mediated sensing guides multisensory grasping as if objects were sensed directly with the hand. This similarity suggests an effective translation of haptic information from the hand operating the tool into object-relevant information for multisensory grasping performance.

The striking resemblance between tool-mediated and hand-based sensing in multisensory grasping is further highlighted by the type of object information integrated with vision: as in hand-based multisensory grasping (Camponogara & Volcic, 2019b, 2021, 2022), tool sensed positional information was sufficient to support vision. The object localization may have occurred by two concurrent processes, consisting of encoding the pattern of somatosensory inputs elicited by the tool’s impact with the object (Miller et al., 2019, 2018) and/or the haptically experienced inertia stemming from the active tool movement, which may have been used to infer the tool’s length and thus the object position at the end of it (Chan, 1994; Solomon & Turvey, 1988; Solomon et al., 1989). Patients deprived of proprioception, indeed, show reduced or even absent effects of action performance when using a tool: the usually observed change in hand kinematics after tool use (Cardinali et al., 2009) fades away when proprioceptive inputs are prevented (Cardinali, Brozzoli, Luauté, Roy, & Farnè, 2016). A reduction in action performance following tool use is also observed when the haptically experienced inertia is prevented by passive tool movements (i.e., tool moved by the experimenter) (Farnè, Iriki, & Làdavas, 2005; Hihara, Obayashi, Tanaka, & Iriki, 2003; lriki, Tanaka, & Iwamura, 1996; Maravita, Spence, Kennett, & Driver, 2002; Obayashi, Tanaka, & Iriki, 2000). Thus, the inherent somatosensory and haptic stimulation characterizing the tool’s active placement on the object may have played a key role in establishing the tool’s length and, consequently, the object’s location.

Even though our results suggest that the tool supports vision by providing mainly positional information, it may be that, in specific visual conditions, the tool-sensed size also plays a role. In haptic-based multisensory grasping (i.e., grasping a seen hand-held object), the haptic size plays only a marginal role in optimal visual conditions (Camponogara & Volcic, 2019b, 2021), whereas it provides a significant contribution to action performance in conditions of visual uncertainty (Camponogara & Volcic, 2022). Thus, introducing visual uncertainty when grasping a tool-held object may lead to a gradual use of the tool-sensed size gathered through the haptically sensed distance between the thumb and the other digits holding the tool’s handle. A second factor, other than optimal vision, that may have prevented the use of tool-sensed size is the unequal mapping between the hand controlling the handle and the aperture of the gripper. When the hand was semi-open or semi-closed, the gripper was either completely open or closed, respectively. This unequal mapping may have prevented the use of the haptic distance between the digits holding the handle to infer the object size (Takahashi et al., 2009; Takahashi & Watt, 2014, 2017).

The successful use of tool-sensed information for action execution hints at common neural structures that govern both tool-mediated and hand-based multisensory grasping. Indeed, the primary motor and somatosensory cortices and the posterior parietal cortex have been shown to play a role in both tool-mediated and hand-based sensing (Gallivan et al., 2013; Jacobs et al., 2010; Johnson-Frey, 2004; Maravita & Iriki, 2004; Miller et al., 2019). These neural structures are also involved in the sensorymotor transformation process of grasping and reaching movements toward visual and haptic targets (Bernier & Grafton, 2010; Buneo, Jarvis, Batista, & Andersen, 2002; Cohen & Andersen, 2002). Thus, transforming haptic information about the tool (i.e., length of the tool, size of the handle) into object positional and size information for action guidance may rely on the same neural circuits involved in processing haptic information (i.e., arm extension and digits separation) stemming from the direct contact of the hand with the object (Berryman et al., 2006; Proske & Gandevia, 2012).

The discovery that a tool can be used as a sensory device to guide a contralateral limb movement has strong practical relevance for prosthesis engineering and rehabilitation studies. It has been shown that the sense of touch could be restored in amputees through special prosthetic devices equipped with microelectrodes, surgically implanted in the amputees’ nerves. During object manipulation, the prosthetic hand movement activates the microelectrodes, which elicit the sensory nerves. The decoding of the nerves activation pattern allows recognizing the size of the held objects to an almost comparable level as in control participants (D’Anna et al., 2019). Our study suggests that amputees could use the restored haptic information to develop and fine-tune bimanual object manipulation skills. Additionally, it also hints at the use of sensory stimulation and multisensory integration techniques to improve prosthesis compliance. Through bimanual object manipulations, prosthesis users could associate the visual with the felt object features (i.e., position and size) and improve prosthesis control. The overlap of neural structures involved in the tool- and hand-mediated sensing can also be successfully exploited to promote the restoration of motor functions following a stroke. By incorporating tasks that require the use of a tool or the hand to guide reaching and grasping movements with their affected hand, stroke patients could practice and improve their motor skills in more varied contexts. This type of interleaved training could promote generalized learning and help maintain the gains made during rehabilitation by improving hand dexterity and coordination.

In conclusion, our study demonstrates that humans can not only use tools as pure grasping or sensory devices, but they can integrate tool-sensed object information with vision to guide fine motor skills, such as precision grips.

## Acknowledgments

We acknowledge the support of the NYU Abu Dhabi Research Enhancement Fund (grant RE183). This work was partially supported by the NYUAD Center for Artificial Intelligence and Robotics, funded by Tamkeen under the NYUAD Research Institute Award CG010.

## Data Availability

Data of all the performed experiments are available at the following link https://osf.io/7n2g5/

